# Discovery of a novel jingmenvirus in Australian sugarcane Soldier fly (*Inopus flavus*) larvae

**DOI:** 10.1101/2022.03.14.484210

**Authors:** Agathe M.G. Colmant, Michael Furlong, Kayvan Etebari

## Abstract

In Australia, soldier flies are major pests of sugarcane, and they can cause significant yield losses in some areas, possibly due to virus transmission to the plants. We sequenced fly larvae salivary glands and identified a novel jingmenvirus, putatively named Inopus flavus jingmenvirus 1 (IFJV1). Phylogenetic trees confirmed that IFJV1 groups with insect-associated jingmenviruses, newly identified flavivirus-like viruses with a segmented genome. After the design and validation of molecular detection systems for IFJV1, larval homogenates were passaged on insect and vertebrate cells but IFJV1 could only be detected in the first two passages in insect cells and not at all in vertebrate cells. Despite this lack of consistent replication in laboratory models, this virus does replicate in its host *Inopus flavus*, as sequenced small RNA from larvae match the IFJV1 sequences, are predominantly 21 nucleotides-long and map to the whole sequences on both strands, which is typical of an actively replicating virus. This discovery confirms the worldwide presence of jingmenviruses, which until now had only been detected on four continents. However, the study of IFJV1 tropism and of the possible pathogenicity to its host or the sugarcane it parasitizes requires the development of a stable replication model.

## Introduction

Sugar cane soldier flies are important pests as they cause significant yield losses in some sugarcane regions in Australia. Soldier flies represent a species complex that comprises at least six endemic species that are economically important pests of sugarcane (1). The damage caused by one of the species, *Inopus flavus* (Diptera: Stratiomyidae), has become more obvious recent years in Australia even though its distribution is believed to be restricted to eastern central Queensland (2,3). Soldier fly pest management is difficult in sugarcane crops as insecticides are ineffective and varietal tolerance to larval feeding is limited. Small numbers of larvae can cause significant damage to the plant and reduce the crop yields. We aimed to investigate whether this effect is linked to soldier fly larvae transmitting viruses to the plant during feeding. Moreover, insect RNA viruses are capable of causing significant reduction in the field populations of agricultural and forestry pests. There are several examples of using insect-specific RNA viruses being used as biological control agents, sometimes in combination with genetically engineered crops (4). In the context of this study, novel viruses are therefore of particular interest due to their potential as plant or insect pathogens, which would then need to be either managed or could be harnessed as biological control agents. The advent of next generation sequencing (NGS) technology has created a great opportunity for novel virus discovery.

Jingmenviruses are a group of novel positive single-stranded RNA (+ssRNA) viruses, currently designated by the ICTV as an unclassified sub-genus in the *Flavivirus* genus, *Flaviviridae* family, which include viruses of medical and veterinary importance such as tick-borne encephalitis or dengue viruses. The first jingmenvirus identified as such was Jingmen tick virus (JMTV), by Qin *et al*. in 2014, in *Rhipicephalus microplus* ticks collected in China (5). JMTV has a four-segmented genome. The first and third segments present one open reading frame (ORF) each, coding for the non-structural proteins 1 and 2 respectively, similar to the NS5 (RNA-dependant-RNA-polymerase (RdRp) and methyltransferase) and NS2B/NS3 (serine protease and helicase) flavivirus proteins. Segments 2 and 4 code for two structural proteins each (VP4 and VP1; VP2 and VP3 respectively) and are more genetically distant from flaviviruses than the non-structural proteins. Since this discovery, several other virus species have been discovered, associated with ticks or human infections (Alongshan virus, Yanggou tick virus, Xinjiang tick virus 1 and Takashi virus), other vertebrates (bats and rodents), insects, or plants (6–11). To date, jingmenviruses have been detected in a wide range of hosts and from geographical locations on four continents (Asia, America, Africa and Europe). Very few reports describe isolation attempts, particularly for the insect-associated viruses, while the replication of the prototype virus JMTV could only be detected for a couple of passages on vertebrate (DH82, dog) and insect (C6/36, mosquito) cell lines or in intracranially injected newborn mice (5).

Identification of new insect viruses and further investigation of their impact on soldier fly populations in different regions will provide a better understanding of the potential interactions between insect specific viruses and their hosts, which could potentially lead to identification of new biological control agents.

## Material and Methods

### Sample collection and RNA extraction

Sugarcane yellow Soldier fly (*Inopus flavus*) larvae were collected from an infested sugarcane field near Hay Point, Queensland (21°18’5”S, 149°14’7”E) in 2019. Sugarcane stools were excavated from the ground and large larvae were manually collected from the roots and associated soil. Larvae were transported to the University of Queensland’s laboratory for RNA extraction and next generation sequencing. The approach was unbiased, since no attempt was made to enrich viral particles through filtration, centrifugation or nuclease treatment. Total RNA samples were extracted from larvae salivary glands as previously described in Etebari *et al*. 2020 (2). Briefly, the larval body surfaces were disinfected by soaking in 75% ethanol for 30 seconds and rinsing in phosphate-buffered saline (PBS). The salivary glands (SG) were then extracted by pulling out the head capsule and removing all other tissues, such as fat body droplets. The SG tissues were pooled and transferred to Qiazol lysis reagent for RNA extraction according to manufacture instruction (QIAGEN; Cat No.: 79306) in pools of 20. Six pools were sent for total RNA sequencing on a HiSeq 4000 (Illumina) by the Australian Genome Research Facility (AGRF, Melbourne), after DNase treatment and an RNA quality control check. Deep sequencing raw data have been deposited in the National Centre for Biotechnology Information’s (NCBI’s) Gene Expression Omnibus (GEO) and are accessible through GEO series accession number GSE127658.

### Transcriptome data analysis and virus discovery

In this study, the CLC Genomics Workbench version 20.0.1 (Qiagen, USA) was used for bioinformatics analyses. All libraries were trimmed from any remaining vector or adapter sequences. The reads were 75bp pair ended with an average fragment size of 350 bp and insert size of 230 bp. Low quality reads (quality score below 0.05) and reads with more than two ambiguous nucleotides were discarded. As the *I. flavus* genome is not sequenced, all reads were mapped to the black soldier fly genome, *Hermetia illucens*, (closest available relative) as proxy genome reference (GCF_905115235.1) to remove insect-related reads. Unmapped reads were retained for *de novo* assembly and virus discovery.

Contigs were constructed with kmer size 45, bubble size 50, and a minimum length of 500 bp, then corrected by mapping all reads against the assembled sequences (min. length fraction=0.9, maximum mismatches=2). The generated contigs were compared to the NCBI viral database using local BLAST and BLASTx algorithms. The e-value was set to 1×10^-10 to maintain high sensitivity and a low false-positive rate. To obtain the segment 2 sequence, the contigs were compared to a database comprising all available sequences from jingmenvirus segments with a glycoprotein (segment 4 for mosquito associated-jingmenviruses Guaico Culex virus and Mole Culex virus, and segment 2 for all other jingmenviruses).

Putative jingmenvirus sequences were re-mapped to RNA-Seq data to inspect for sufficient coverage and possible mis-assembly. The CLC Genomic Workbench’s RNA-Seq function (min. length fraction=0.9, maximum mismatches=2, insertion cost=3, deletion cost=3) on a non-strand specific option was used.

The signal peptide, potential glycosylation sites and transmembrane domains were predicted using the online tools SignalP 6.0 (https://services.healthtech.dtu.dk/service.php?SignalP), NetNGlyc (https://services.healthtech.dtu.dk/service.php?NetNGlyc-1.0) and TMHMM (https://services.healthtech.dtu.dk/service.php?TMHMM-2.0) (12).

### Phylogenetic analysis

The NSP1 and NSP2 amino acid (aa) sequence of both tick- and insect-associated jingmenviruses including those of IFJV1 were used to build phylogenetic trees. First, multiple aa sequence alignments were performed with MUSCLE using Geneious Prime. Then, the maximum-likelihood phylogenetic trees were inferred using a JTT substitution matrix and assuming a discretised gamma rate distribution with four rate categories (shape parameter fixed at 1.0) and with 1000 bootstraps.

### In vitro *isolation attempts*

The samples used for the isolation attempts were whole larval body homogenates corresponding to the positive salivary glands, homogenised in 500μL phosphate buffered saline (PBS, pH 7.4) with or without the addition of phenylthiourea (PTU) at saturation, to prevent melanisation.

The cell lines used for the isolation attempts were C6/36 *Aedes albopictus*-derived cells, maintained at 28°C in RPMI 1640 with 2-5% fetal bovine serum (FBS); S2 *Drosophila melanogaster*-derived cells maintained at 28°C in Schneider’s drosophila medium with 10% FBS; BSR baby hamster kidney *Mesocricetus auratus*-derived and Vero African green monkey kidney *Cercopithecus aethiops*-derived cells, maintained at 37°C with 5% CO2 in DMEM with 5% FBS. All cell culture media were supplemented with 50 U/mL of penicillin, 50 μg/mL of streptomycin, 2 mmol/L of L-glutamine.

Cells were seeded with 1×10^5 cells per well in 24-well plates and incubated overnight at 28°C or 37°C for insect and vertebrate cells respectively. The cell culture supernatant was removed and replaced with 200µL inoculum (homogenate diluted 1/10, filtered through a 0.22 µm sterile filter). The cells were left to incubate at room temperature for 30 minutes on a rocker and a further 60 minutes at 28°C (insect) or 37°C (vertebrate). After incubation, the inoculum was removed, 50µL were reserved for RNA extraction, and the cells were washed three times with 750uL sterile phosphate buffered saline (PBS). The cell culture medium was replenished with 750µL fresh medium with 2% FBS. These cultures were left to incubate for 7 days at 28°C or 37°C for insect and vertebrate cells respectively. After incubation, RNA was extracted from the harvested supernatants, which were also passaged on freshly seeded cells. RNA extractions were performed using the Machery Nagel Nucleospin RNA extraction kit following the manufacturer’s instructions. The RNA extracts were used as templates with the Superscript III One-Step RT-PCR System with Platinum Taq DNA polymerase kit (Invitrogen) following the manufacturer’s instructions and primers designed to detect the jingmenvirus segments, see Table 1.

**Table 1.**
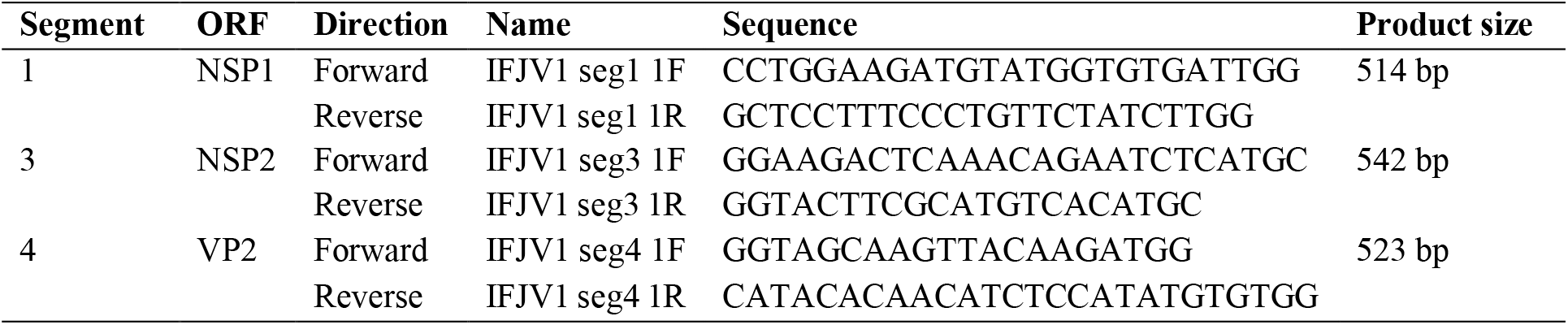
Primers designed to detect IFJV1

### Viral derived small RNA analysis

To analyse the host RNAi response to IFJV1 and determine whether the virus was replicating in its host, a small RNA (sRNA) library was generated from one of the positive pools of 20 SG from starved individuals using the NEBNext® Multiplex Small RNA Library Prep Kit for Illumina® at the Novogene Genomics Singapore Pte Ltd. The purified cDNA libraries were sequenced on Novaseq 6000 (SE50) and raw sequencing reads were obtained using Illumina’s Sequencing Control Studio software. Raw data were stripped of adapters and reads with a quality score above 0.05, and less than two ambiguous nucleotides were retained. Reads without 3’ adapters and reads with less than 16 nt were discarded. The clean reads were mapped to each IFJV1segment. We examined both the size distribution of the viral-derived sRNA fragments as well as the genomic distribution for each segment on both strands (positive- and negative-sense).

## Results

### Identification of Inopus flavus jingmenvirus 1

We identified several RNA viruses from the RNAseq libraries of salivary glands from 6 pools of 20 SG from soldier fly larvae, collected from north Queensland, Australia, in 2019. Here, we will focus only on a novel jingmenvirus we discovered and putatively named Inopus flavus jingmenvirus 1 (IFJV1). The sequences obtained for IFJV1 cover the putative full coding sequences of the four segments. The organisation of the IFJV1 genome follows what has been found previously for insect-associated jingmenviruses (Figure 1). Segment 1 codes for NSP1, a 924 aa-long non-structural protein with RdRp and methyltransferase domains. Segment 2 is bicistronic and codes for the 485 aa-long glycoprotein VP1 with four transmembrane domains and the putative 104 aa-long structural protein VP4 with two transmembrane domains. Segment 3 codes for the 805 aa-long second non-structural protein, NSP2, which contains serine protease and helicase domains. Segment 4 is bicistronic and codes for two 254 and 471 aa-long structural proteins VP2 and VP3. VP2 has a signal peptide with a predicted cleavage site between positions 16 and 17 and no predicted N-glycosylation sites, while VP3 contains at least six transmembrane domains. These proteins are the putative equivalent of the capsid and membrane proteins found in flaviviruses. The IFJV1 genomic sequences have been deposited on Genbank and have been assigned the following accession numbers: OM869459-OM869462.

**Figure 1.**
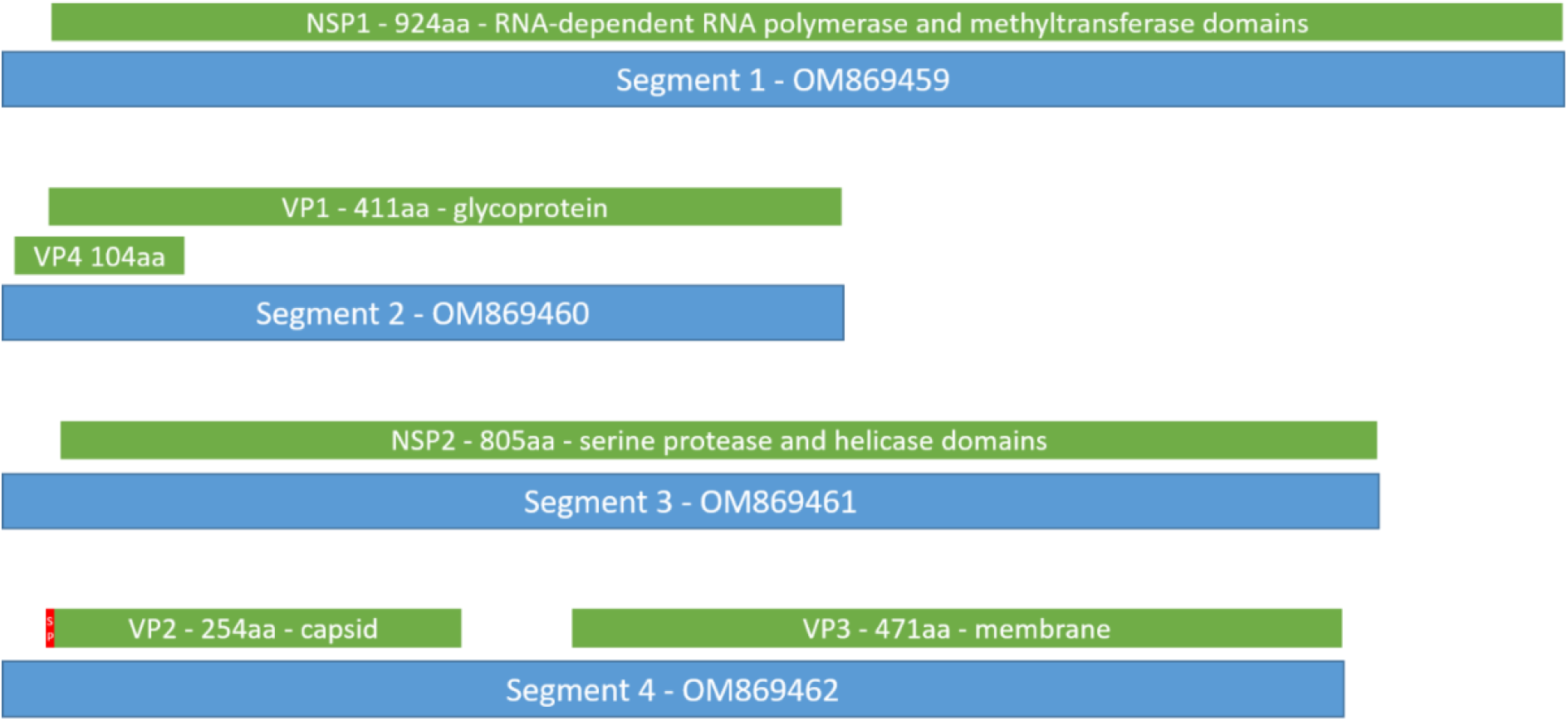
Genome organisation of IFJV1. Segments are in blue, open reading frames are in green and the signal peptide in red. SP: signal peptide; aa: amino acid. Genbank accession numbers OM869459-OM869462.

### Phylogenetic analysis

The ORF aa sequences were compared to published sequences using NCBI BLAST, with the blastp algorithm. According to these comparisons, IFJV1 is most similar to Mole Culex virus and Guaico Culex virus, two insect-associated jingmenviruses isolated from mosquitoes **(13,14)**. Their NSP1 share around 50% aa identity, the structural proteins VP2 and VP3 share approximately 25% aa identity and the NSP2 share 40% aa identity. These identity percentages leave no doubt that IFJV1 is indeed a novel species of virus. The clustering of IFJV1 sequences with the mosquito-associated jingmenviruses within the insect-associated clade of jingmenviruses was confirmed by phylogenetic analysis on the aa sequences of NSP1 and NSP2 (Figure 2).

**Figure 2.**
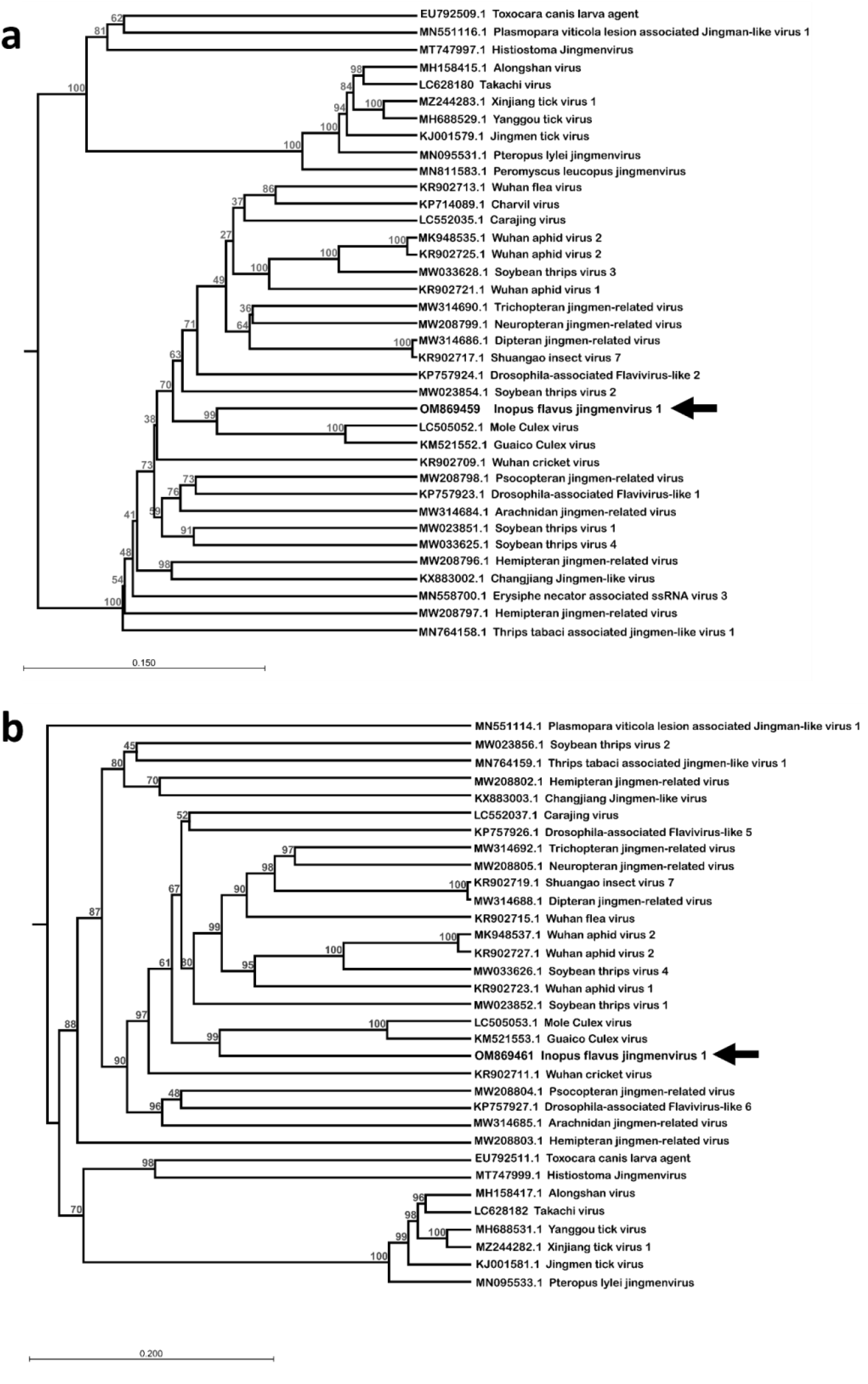
Phylogenetic analysis of the amino acid sequence of NSP1 (A) and NSP2 (B) of Inopus flavus jingmenvirus 1 aligned with reference jingmenviruses. These maximum-likelihood phylogenies were inferred using a JTT substitution matrix and assuming a discretised gamma rate distribution with four rate categories and with 100 bootstraps. Bar: branch length.

### In vitro *virus isolation*

Two homogenates from the bodies of soldier fly larvae with IFJV1 positive salivary glands were passaged on insect cell lines, C6/36 mosquito- and S2 drosophila-derived cells in an attempt to isolate the virus. While sequences from the three known segments were detected in the inoculum, the signal was lost at the third passage, suggesting that the cells were not able to stably support the viral replication. A similar attempt was made on vertebrate BSR hamster- and Vero monkey-derived cells, but no IFJV1 replication was detected in these cells either.

### Virus derived small RNA profiles

To determine if the virus was replicating in its soldier fly host, despite its inability to replicate in classical *in vitro* laboratory models, we analysed the sRNA in one of the positive pools of 20 SG from starved larvae. During virus infection in arthropods, virus-related double-stranded RNA (dsRNA) triggers the activity of host RNAi responses and the host riboendonuclease III enzyme Dicer-2 cleaves this dsRNA into virus-derived small interfering RNAs (vsiRNAs) which are 19–22 nt in length (15). These vsiRNAs are then loaded into the RNA-induced silencing complex, where they target RNA molecules through complementarity, reduce the virus gene transcription and ultimately virus replication. For insect viruses the vsiRNAs display a sharp peak in 21 nt, are symmetrically distributed throughout the viral genome, and map to both strands (positive and negative) (16–19). Therefore, after selecting only the reads between 18 and 30 nt in length, we mapped the sRNA libraries to the three IFJV1 segment sequences and generated the size distribution graphs (Figure 3). We found that the virus-derived sRNAs displayed a peak in size at 21 nt in length, and when the 21 nt reads were mapped back to each segment, we observed that most of the four sequences were covered by the sRNA reads, on both strands. This suggests that IFJV1 is replicating actively in its soldier fly host, and activates its immune response.

**Figure 3.**
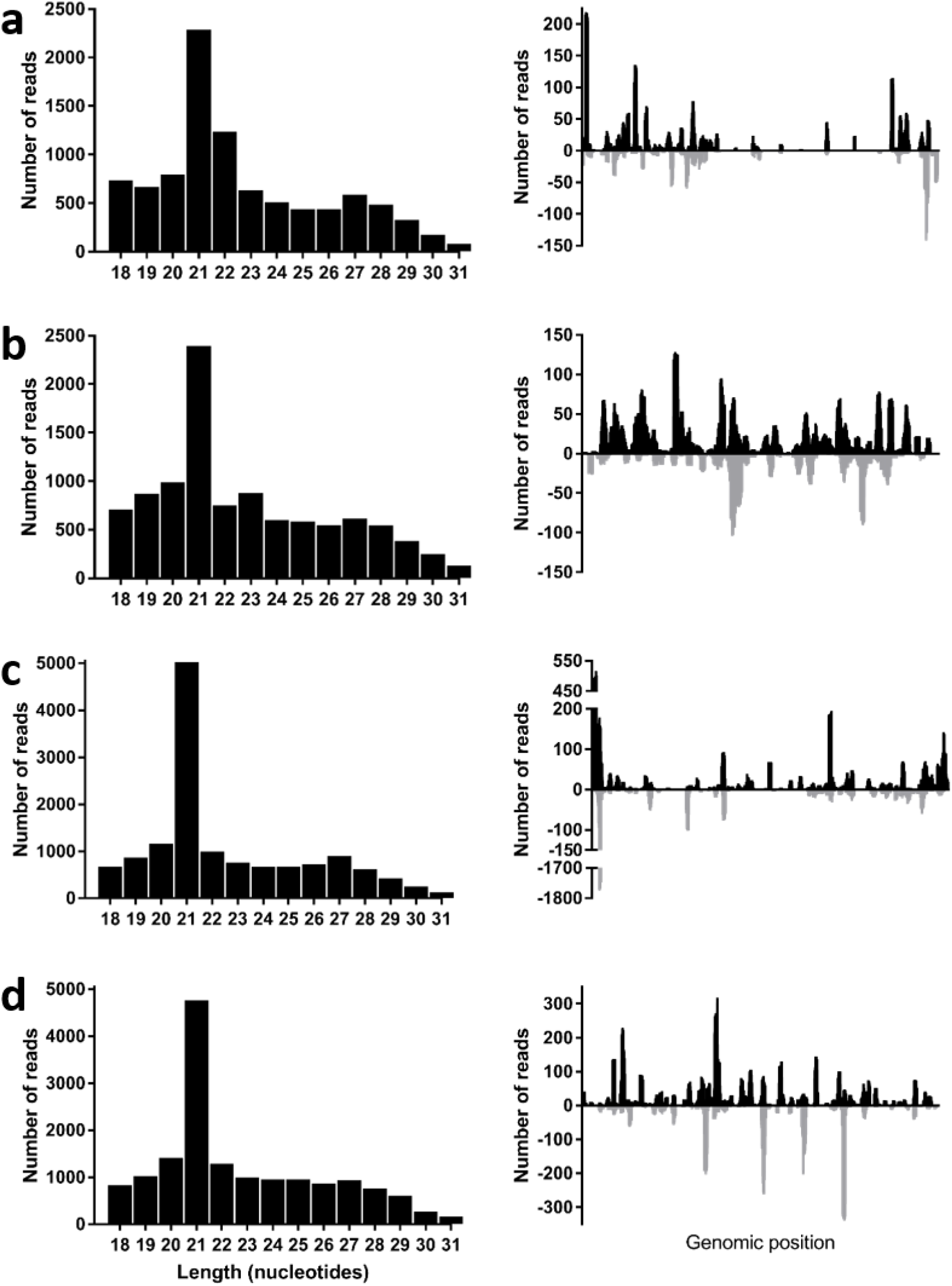
Length profile and distribution of IFJV1-derived sRNAs. The virus-derived sRNA length profiles are on the left (A: segment 1; B: segment 2, C: segment 3; D: segment 4) and the distribution of 21 nt-long IFJV1-derived sRNA mapped back to the virus positive (black) and negative (grey) sense nucleotide sequences are on the right.

## Discussion

Here we report the discovery and identification of a novel jingmenvirus in Australian sugarcane soldier flies. This is the first report of a jingmenvirus in Oceania, which extends the distribution of jingmenviruses to all continents except Antarctica. This ubiquitous distribution highlights that jingmenviruses should be studied and their emerging potential characterised. Moreover, the close association between *I. flavus* larvae (the stage in which the virus was found) and sugar cane leads us to question whether the virus replicates within the plants and/or interacts with it in another way. This consideration is all the more relevant, given that Wuhan aphid virus 2 was detected both in an insect (*Hyalopterus pruni*) and in a plant (*Pisum sativum*), with the two strains displaying high levels of sequence similarity **(20,21)**. In order to investigate the effect of the virus in *I. flavus* and its potential as a biological control agent of the pest, its pathogenicity, prevalence and effect on *I. flavus* populations need to be understood, alongside its potential for interference with viruses co-circulating in soldier flies and sugar cane.

Interestingly, the IFJV1 sequences clustered with the two mosquito-associated jingmenviruses, Guaico Culex virus and Mole Culex virus **(13,14)**. However, while mosquitoes and soldier flies are both belong to the *Diptera*, other jingmenviruses that were detected in dipteran insects such as Drosophilae or Culicoides do not cluster together. Furthermore, the jingmenviruses that have been detected in hemipteran hosts do not cluster together, nor with those found in the closely related families in order *Thysanoptera* and *Psocodea* **(22)**. The phylogenetic organisation of insect-related jingmenviruses therefore does not follow the phylogenetic organisation of their identified hosts **(22)**. This observation is corroborated by the presence within the phylogeny of jingmenviruses detected in non-insect arthropods from two other classes, Louisiana crawfish (*Procambarus clarkia;* Malacostraca) and small wood scorpions (*Euscorpius sicanus;* Arachnida) as well as organisms from the Fungal (*Erysiphe necator* and *Plasmopara viticola*) and Plant (*Pisum sativum*) kingdoms **(21,23–26)**. This lack of correlation between the virus and host phylogenies suggests that the viruses have not co-evolved with their hosts. Since most of the jingmenvirus sequences originate from metagenomics studies, this observed phenomenon could be due to an incorrect host assignment, since the method would not differentiate sequences from an insect or from its previous meal, or a contaminating parasite or fungus.

Another reason for the observed discrepancy between host and virus phylogenies could be the existence of a reservoir for these viruses, outside of the *Insecta* class, for example in plants or fungi as suggested above. The viruses could be transmitted to the insect hosts from this reservoir, enabling independent evolution. In any case, jingmenviruses need to be studied further to elucidate their ecology and modes of transmission. Unfortunately, we were not able to further investigate these considerations, since the virus could not be cultured stably *in vitro* in any cell line. While this has been found for other jingmenviruses, the mechanisms involved in this replication restriction in laboratory models are not yet clear (5,27,28). We have shown that IFJV1 can elicit an immune response from its host *via* the RNAi pathways by analyzing the sRNA in *Inopus flavus* homogenates, but the C6/36 cell line has a dysfunctional RNAi response, so this mechanism is unlikely to prevent IFJV1 replication *in vitro* in these cells (29). The presence of a IFJV1 specific sRNA response is however proof that the virus is replication-competent in its host, since vsiRNAs are produced in the presence of dsRNA, which only occurs in the form of replicative intermediates for +ssRNA viruses. These data therefore show that the lack of replication observed *in vitro* is due to an inappropriate model rather than a replication-incompetent virus. The sRNA data also demonstrate that IFJV1 is indeed an *I. flavus* virus and that this host has been assigned correctly.

Overall, our study increases the knowledge on jingmenviruses by adding a new member from a new continent and new host to this sub-genus. This report also demonstrates the importance of developing a stable laboratory model for the replication of jingmenviruses, for their thorough characterization and the evaluation of their potential to emerge as insect, plant or vertebrate pathogens.

## Acknowledgements

This project was supported by Sugar Research Australia funding (SRA-00504). We would also like to thank Karel. R. Lindsay and Manda Khudhir for assistance in collecting soldier fly larvae.

## Authors’ contribution

**Agathe M.G. Colmant:** Methodology, Formal analysis, Investigation, Data Curation, Visualization, Writing - Original Draft, Writing - Review & Editing

**Michael Furlong:** Investigation, Resources, Project administration, Funding acquisition, Writing - Review & Editing

**Kayvan Etebari:** Conceptualization, Methodology, Formal analysis, Investigation, Data Curation, Visualization, Writing - Review & Editing

